# Generation of Aggregates of Mouse ES Cells that Show Symmetry Breaking, Polarisation and Emergent Collective Behaviour *in vitro*

**DOI:** 10.1101/005215

**Authors:** Peter Baillie-Johnson, Susanne C. van den Brink, Tina Balayo, David A. Turner, Alfonso Martinez Arias

## Abstract

Dissociated mouse embryonic stem (ES) cells were cultured to form aggregates in small volumes of basal medium in U-bottomed, non tissue-culture-treated 96-well plates and subsequently maintained in suspension culture. After growth for 48 hours, the aggregates are competent to respond to ubiquitous experimental signals which result in their symmetry-breaking and generation of defined polarised structures by 96 hours. It is envisaged that this system can be applied both to the study of early developmental events and more broadly to the processes of self-organisation and cellular decision-making. It may also provide a suitable niche for the generation of cell types present in the embryo but unobtainable from conventional adherent culture.

## Introduction

The study and understanding of cell-fate decisions in early mammalian development can make use of cultures of Embryonic Stem Cells (ESCs), clonal populations derived from blastocysts which have the ability to self renew and differentiate into all cell types of an organism i.e. they are pluripotent (Evans and Kaufman, 1981; Martin, 1981). While these cultures have been and are continuing to be useful for the understanding of the molecular basis of cell-fate decisions, they are unable to reproduce some of the spatial arrangements that are generated in embryos. Thus, for example, in the embryo the process of gastrulation transforms a single epithelial layer into the three distinct germ layers and endows the embryo with an overt anterioposterior organisation (Ramkumar and Anderson, 2011; Solnica-Krezel and Sepich, 2012; Tam and Gad, 2004). Attempts to recapitulate these events *ex vivo* have been based on the generation of three-dimensional aggregates of ESCs, referred to as embryoid bodies (EBs), subjecting them to differentiation conditions. These aggregates can be coaxed to differentiate into many different cell types, some of them not efficiently induced in adherent culture e.g. blood (Nostro et al., 2008). However they are unable to display the morphogenetic behaviour, germ layer distribution or axial organization that are characteristics of embryos, resulting in spatial disorganization (Leahy et al., 1999; Sajini et al., 2012). In one report, treatment of EBs with Wnt leads to a weak polarization in gene expression in some aggregates but no clear morphogenesis is observed (Berge et al., 2008). In more recent reports, EBs that have been cultured for extended periods of time develop anterior structures, retinas and cortex, which mimic their embryonic counterparts but develop without the context of an axial organization (Eiraku et al., 2011; Lancaster et al., 2013).

Working with aggregates of mouse P19 Embryo Carcinoma (EC) cells, Marikawa *et al*. (Marikawa et al., 2009) have reported the emergence of elongated structures of mesodermal origin reminiscent of the elongations that are observed with exogastrulae in amphibian and sea urchin embryos and Keller explants (Holtfreter, 1933; Horstadius, 1939; Ishihara et al., 1982; Keller and Danilchik, 1988). This observation led us to attempt to reproduce the behaviour of the P19 EC cells. Here we report culture conditions that lead to symmetry breaking and axial organization in aggregates of ESCs. The protocol was inspired by the work of Marikawa (Marikawa et al., 2009) but was progressively modified to optimize the behaviour of the cells in the direction of their behaviour in embryos (our protocol is described in detail below). An important difference from the work with P19 cells is that the aggregation was performed in a 96-well plate instead of hanging drops in the manner of Eiraku *et al*. (Eiraku et al., 2008) who reported improved efficiency and throughput from a Serum-Free culture of Embryoid Body-like aggregates in these wells (SFEBq method). This change resulted in an increased efficiency of the method. An additional and significant difference between this method and the hanging drops is that the SFEBq method maintains the aggregates in individual culture, thereby eliminating fusions that occur when pooling aggregates from hanging drops. Further improvements are described and discussed in the detailed protocol below.

### Applications of the Method

The aggregation system described here leads to symmetry breaking events, axial elongation, germ layer specification and gastrulation like movements (Turner et al., 2014; van den Brink et al., 2014). For these reasons the method has the potential to allow detailed mechanistic analysis of processes that may be difficult to study in the embryo. Furthermore, a potential application is in the generation of tissues and organs that are not easily obtainable in adherent culture ((Turner et al., 2014) submitted). Three-dimensional aggregate culture offers a physical structure and signalling environment that may not be achievable by conventional means, leading to a new approach for the derivation of embryonic lineages.

### Comparison with Other Methods

Our method aims to generate axial derivatives in culture and this contrasts with existing methods for the generation of organs from ES cells that lead to the emergence of optic cup (Eiraku et al., 2011) and cerebral cortex development (Lancaster et al., 2013). Both systems use a short period of aggregation in low-adhesion 96-well plates before aggregated cells are transferred to Matrigel droplets for on-going culture. Eiraku *et al*. describe “fully stratified neural retina tissue architecture in this ES cell culture self-forms in a spatiotemporally regulated manner mimicking *in vivo* development” (Eiraku et al., 2011). The culture system aggregates approximately 3000 mouse ES cells in a 100µl volume according to the SFEBq method described previously. Matrigel is applied at the end of the first day and the aggregates are transferred to suspension culture for a further 8-11 days in 40% O_2_ and 5% CO_2_. By contrast, Lancaster *et al*. aggregated human ES cells at a high density (4500 cells per aggregate) in a low-adhesion 96-well plate for 6 days, before transfer to a low-adhesion 24-well plate for a further 5 days. These aggregates, which Lancaster *et al*. term *cerebral organoids* were transferred to Matrigel on the 11^th^ day of culture and maintained in stationary culture for a further 4 days, before they were transferred to a spinning bioreactor for on-going culture. Analysis of the cerebral organoids found them to have developed a number of regions similar in morphology to brain (Lancaster et al., 2013). From the development of neural identity (8-10 days) to acquisition of defined brain regions (>20-30 days), this method is comparatively long in duration with respect to the method described by Eiraku *et al*. (Eiraku et al., 2011).

In this manuscript, we have developed a protocol that allows the formation of aggregated mouse ES cells that display overt regional organisation in terms of gene expression, morphology and appearance on a significantly shorter time-scale of up to 5 days (**Fig. 1**). These aggregates, or *gastruloids* (referred to as such due to their ability to display many of the features associated with the gastrulating mouse embryo (van den Brink et al., 2014)) require approximately 5-10 fold fewer cells than those detailed above (a density of which is critical for proper organisation (van den Brink et al., 2014)), and their self-organisation occurs on a global scale rather than the regional examples reported in the studies above.

**Figure 1.**
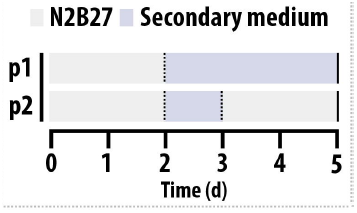
Schematic representation of the differentiation time-courses. Small volumes (see materials and methods) of N2B27 containing a determined concentration of mouse ES cells in suspension are pipetted into the wells of a 96-well plate and left for 2 days to allow their condensation and aggregation (p1, p2). After 2 days (see Fig. 2 centre panel and Fig. 3) when small aggregates have formed, the medium is removed and a secondary medium is added either for the rest of the experiment (p1) or for one day only before being changed back to N2B27 (p2). A pulse of Chi between days 2 and 3 results in efficient and consistent elongated aggregates.

By contrast to the methods described above by Lancaster *et al*. and Eiraku *et al*. (Eiraku et al., 2011; Lancaster et al., 2013), we do not require the transfer of aggregated cells to an artificial matrix, but grow cells in suspension within a neutral, serum free medium (N2B27) in non-adherent, bacterial-grade 96-well plate for 2 days before addition of a defined secondary medium. This 2 day incubation period allows cells to reach a postimplantation epiblast-like state in which they are competent to respond to signals from the secondary medium (Turner et al., 2014; Turner et al., 2013; van den Brink et al., 2014). Characterisation by live imaging or immunocytochemical analysis of the aggregates that develop occurs on day 5, by which time the aggregates have often developed significant changes in morphology and gene expression (**Fig. 2**). Whereas the existing system was developed to focus on the generation of anterior structures, the protocol described in this manuscript allows us to investigate the generation of axial structures and posterior development (Turner et al., 2014; van den Brink et al., 2014).

**Figure 2.**
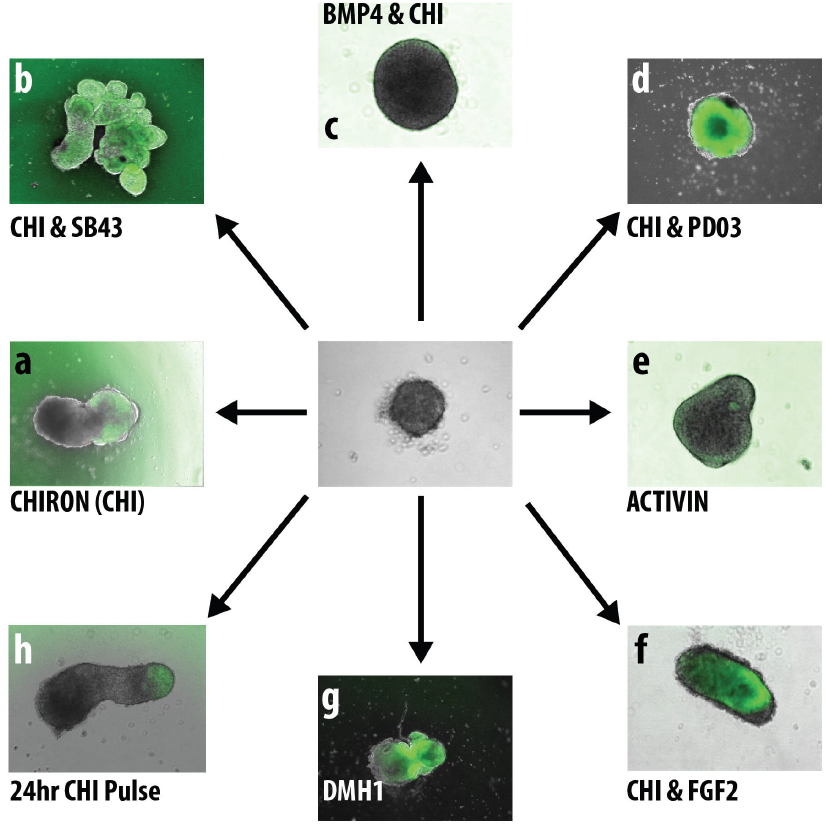
The effect of different treatments on the morphology of aggregated ES cells. Following 2 days in N2B27, ES cells have formed aggregates. Addition of specific factors either continuously (a-e, g) or as a one day pulse between days 2 and 3 (f, h) can alter the phenotype of the aggregates with respect to the polarity of gene expression, elongation potential, or their overall shape. The examples in this figure are aggregates formed from Sox1::GFP mouse ES cells (Ying et al., 2003). See main text for explanation of abbreviations.

### Experimental Design

From previous experimentation, we determined that cells, either in monolayer (Turner et al., 2013) or in an aggregated form (van den Brink et al., 2014) have an optimal time-window in which they can respond to signals promoting the self-organisation and elongation of the aggregated cells. Following the 2 day aggregation period in suspension in N2B27 (**Fig. 1**), we find that a single 24h pulse of a secondary medium containing Chiron (an activator of ß-Catenin signalling) rather than continuous treatment is the optimum condition for generating elongated and organised structures (**Fig. 1**, protocol p2) (Turner et al., 2014).

### Level of Expertise Required

In addition to good, standard tissue-culture practice, accurate cell counting and pipetting, and careful handling of cell suspensions during their initial plating, a minimal level of expertise is required for this technique.

### Limitations

As mentioned previously (and below), it is essential to ensure that the aggregates are maintained in suspension culture and are prevented from adhering to any surface. It is therefore an elementary principle that during the growth of these aggregates, they can only be cultured for approximately 5-6 days post plating until their size renders them competent under gravity to sink to the bottom of the wells and force an adhesion even on non-coated plasticware. At present, this limits the duration of 3D culture but it is likely that the application of artificial matrices in the future will extend this time considerably.

Suspension culture also constrains imaging methods since the aggregates are not immobilised, however modification of imaging systems to allow the imaging of 96-well plates minimises movement and allows effective imaging (van den Brink et al., 2014).

In order to change the medium without removing the aggregates, a small volume must be left in the bottom of the well. The consequences for chemical genetics approaches are the need to consider this dilution and any residual effects from previous media. Current work has not corrected for dilution and it is assumed that daily medium changes will minimise residual effects.

## Materials

### Reagents

0.1% Gelatin in PBS, 0.25% Trypsin-EDTA (Gibco, UK), GMEM (Gibco, UK) supplemented with 10% Foetal Bovine Serum (BioSera) and LIF (hereon referred to as ES+LIF), N2B27 (NDiff 227, StemCells Inc., UK), Activin A (30ng/ml), CHIR99021 (Chi, 3µM; Tocris Biosciences), SB431542 (SB43; 10µM, Tocris Biosciences), BMP4 (1ng/ml), FGF2 (2.5ng/ml), PD0325901 (PD03; 1µM, Tocris Biosciences) and Dorsomorphin-H1 (DMH1, 0.5µM, Tocris Biosciences).

### Equipment

Improved Neubauer haemocytometer or equivalent counting device, 8-channel 200µl pipette, waterbath set to maintain 37°C, humidified and atmospherically maintained sterile tissue-culture incubator (37°C, 5% CO_2_), laminar flow hood, 25 cm^2^ tissue-culture treated culture flasks (T25; for maintenance of cell lines). Sterile and clear U-bottomed 96-well plate for suspension culture (Grenier Bio-One 650185) and sterile plastic reservoirs.

### Reagent Setup

All media should be warmed prior to use in covered (but not sealed) vessels in the incubator to allow temperature and pH equilibration before use and tissue-culture flasks for adherent culture should be coated in 0.1% gelatin for at least 20 minutes prior to passaging. Ensure that any medium that is warmed from frozen is properly mixed before use.

## Procedure

### Plating out

1. Grow mouse ES cells in a gelatin-coated 25cm^2^ tissue-culture treated flask until approximately 80% confluent. Remove spent medium from the stock flask and wash twice with 5ml PBS *Important! Grow for at least two passages post-thawing before beginning this protocol*.
2. Aspirate PBS and add 1ml Trypsin. Leave in incubator for around 2 minutes or until the cells have lifted from the tissue-culture flask.
3. Check for detachment under the microscope, striking the wall of the flask if necessary.
4. Pipette up and down with a P1000 to generate single cell suspension. *Important! A single cell suspension is required for accurate counting and uniform aggregate size*.
5. Neutralise Trypsin with 5ml ES+LIF, washing down the growth surface to maximise cell recovery.
6. Remove a 1ml aliquot and count the cells with the Improved Neubauer haemocytometer.
7. Determine the volume of suspension required to give 10 cells/µl in the final volume: ((Required cell number) / (average cell count x 10 000)) x 1000 = volume required, in microlitres. *e.g. 50 000 cells needed for a 5ml suspension, enough for one 96-well plate*.
8. Add this volume to 5ml pre-warmed PBS in a 15ml Falcon tube.
9. Centrifuge both tubes down at ∼170 RCF for 5 minutes.
10. Aspirate the PBS, being careful not to disturb the pellet.
11. Add 5ml warm PBS to resuspend the pellet and centrifuge as before.
12. Aspirate the medium from the stock tube and resuspend in fresh ES+LIF. Passage to the gelatinised flask as required.
13. Aspirate the PBS from the pellet, again making sure not to agitate it.
14. Resuspend the pellet in 1ml warm N2B27 with a P1000. Make up to the final volume with the remainder of the N2B27.
15. Transfer the suspension to the sterile reservoir and use the multichannel pipette to plate out a 40µl drop into each well of the 96-well plate.
16. Cover the 96-well plate and confirm the presence of cells with an inverted bench-top microscope (Fig. 3A).
17. Leave undisturbed to grow for 2 days.

### Changing to Experimental Media

1. For the first change, add 150µl secondary-medium to each well to the existing 40µl of N2B27. For the generation of highly reproducible, organised, elongated aggregates, a 24h pulse of N2B27 containing 3µM Chi is required after the first 48h N2B27 (Fig. 1 (protocol p2), Fig. 2h).
2. Subsequent changes (daily): gently draw off 150µl from the side of the wells with the multichannel pipette by holding it at an angle. The aggregate should remain in 40µl in the bottom of the well. If pulsed treatment of factors occurred (above, point 1), N2B27 is used for this and subsequent medium changes.
3. Replace with the same volume of warm medium. The force should be sufficient to keep the aggregates in suspension and prevent them adhering to the bottom of the well.

### Removing Aggregates

1. Set a P1000 to ∼200µl and cut approximately 5mm from the end of the tip with a pair of sterile scissors.
2. Draw the medium within the 96-well plate a short way into the tip, expel it to agitate the aggregate, then take up the whole volume.
3. Collect by dispensing into a small volume of pre-warmed medium or PBS in a suitable non-treated vessel. *Large numbers of aggregates can be picked in this way without damage. If the culture is to continue in the plate, ensure the empty wells are refilled with 200µl warm PBS to minimise evaporation*.

## Timing

Preparing gastruloids from a single cell line – 1hour.

Hands-off time while the cells aggregate – 2 days.

Subsequent medium changes – 10 minutes per plate.

Picking aggregates – 15 minutes per plate.

## Troubleshooting

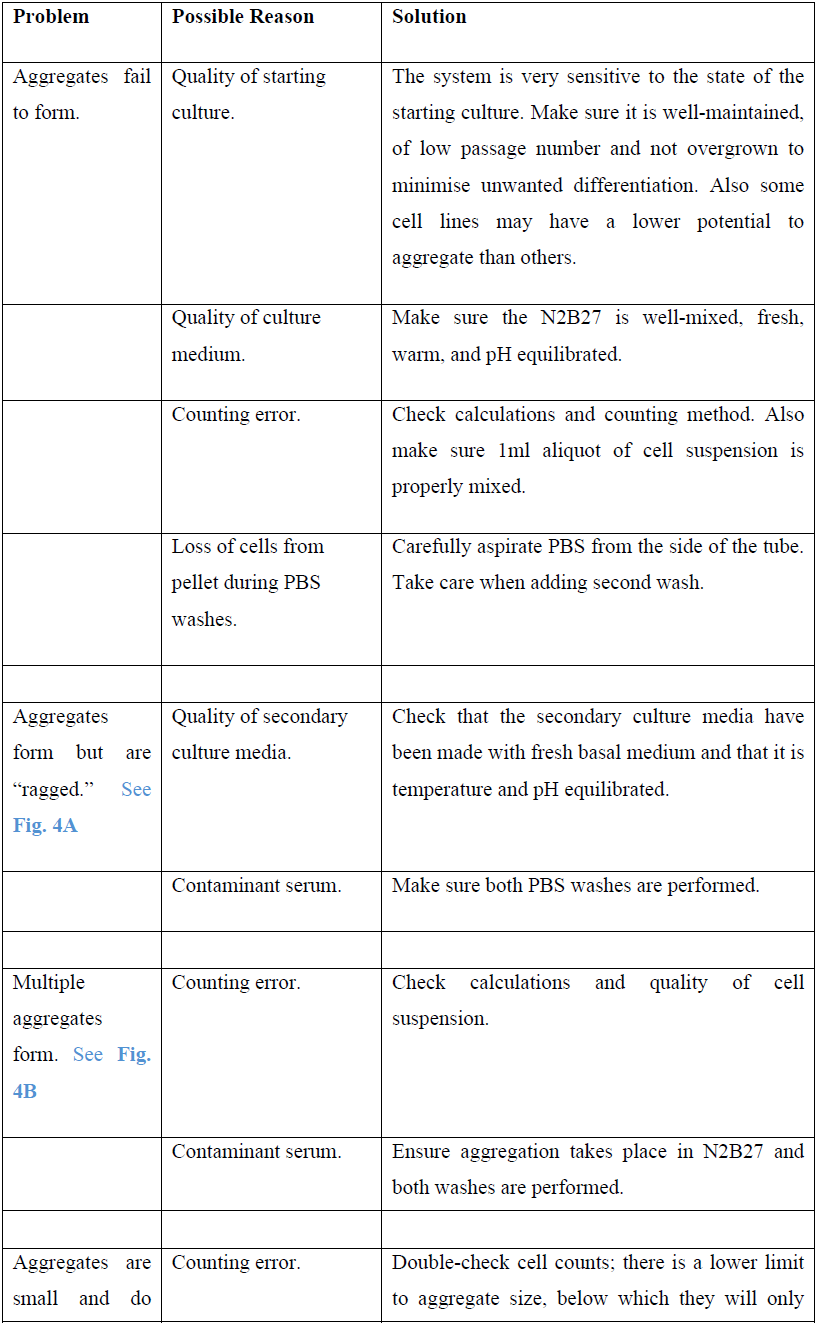
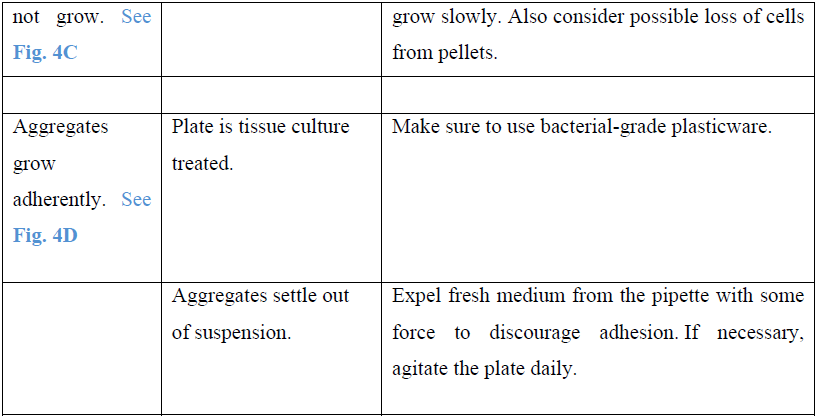

## ANTICIPATED RESULTS

At the end of the first day in N2B27, the cells should have begun to aggregate and compact; individual cells become indistinct (**Fig. 3B**). By the end of the second day, the aggregates should look “clean” and have taken up all the cells (**Fig. 3C**).

**Figure. 3.**
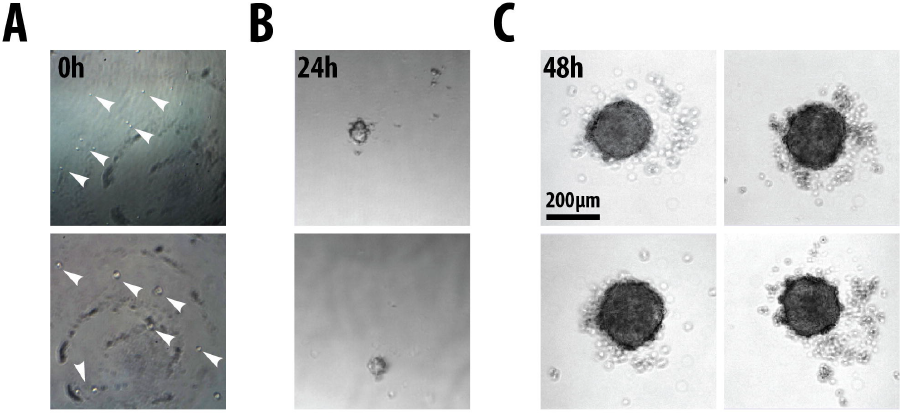
Development of aggregates from single-cell suspensions. (A) Two examples of single-cell suspensions immediately after plating. White arrow-heads indicate selected single-cells. (B) Following 24h culture single-cells have condensed and formed aggregates. Live imaging of the formation of the aggregates in real-time will be available elsewhere ((van den Brink et al., 2014) in preparation). (C) The final day of N2B27 results in larger aggregates without any form of polarisation or elongation. Scale-bar represents 200µm.

**Figure 4.**
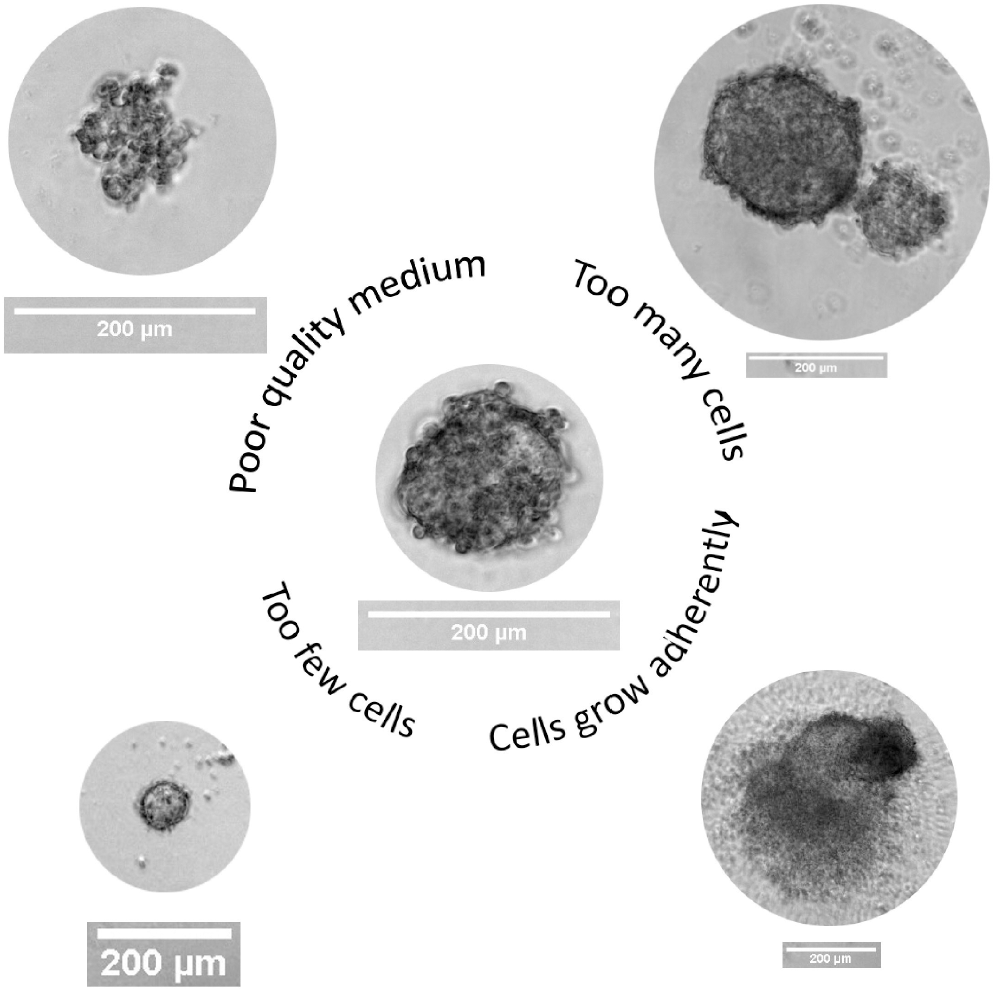
Examples of failures in aggregate formation. The ability to generate reproducible aggregates depends on critical factors such as (A) the quality of secondary medium, (B, C) the accuracy in counting the initial number of cells and (D) and ensuring the aggregates do not form adherent colonies. Typical examples of aggregate formation errors for each of the mentioned conditions (A-D) are shown. Scale-bare represents 200μm; see trouble-shooting table for details.

During the third day of culture, the aggregates are in the secondary media. They will begin to shed cells and respond to these signals between the end of this day, and 12 hours into the fourth day. These responses can be seen as changes in gene expression and morphology, both of which are dependent on the treatment the aggregates have received following the initial 48h aggregation period in N2B27 (**Fig. 2**).

The optimum time for imaging is the end of the fourth day, by which point the aggregates will have developed clear morphologies but should not have begun to settle out of suspension. They can be imaged with widefield microscopy, or fixed and immunostained for confocal work. Early embryo staining protocols are recommended for sufficient antibody penetration.

Under optimum conditions (24h pulse of 3µM Chi on day 2-3; **Fig. 1**, p2 and **Fig. 2h**), aggregates will reliably form in all wells and show clear responses to the experimental manipulations by the fourth day.

## EXAMPLES

In addition to the aggregates shown in **Fig. 2**, further examples of well-formed aggregates at various stages of their development and differentiation will be seen in our manuscripts that are currently in preparation or in submission (Turner et al., 2014; van den Brink et al., 2014).

## Abbreviations

Chi: CHIR99021
DMH1: Dorsomorphin-H1
ES cell: Embryonic Stem Cell
PD03: PD0325901
SB43: SB431542

## Acknowledgements

This work is funded by an ERC Advanced Investigator Award to AMA (DAT, TB) with a contribution of a Project Grant from the Wellcome Trust to AMA, an EPSRC studentship to PB-J and Erasmus, Stichting dr. Hendrik Muller’s Vaderlandsch Fonds and Fundatie van de Vrijvrouwe van Renswoude te ’s-Gravenhageto to SCvdB. We want to thank J. Briscoe, S. Muñoz-Descalzo, C. Schröeter, and T. Rodriguez, for discussions and constructive criticisms.

## Author Contributions

PB-J, SCvdB, TB & DAT optimised and performed the experiments; AMA conceived the project and PB-J, AMA & DAT wrote the manuscript. All authors read and approved the final manuscript.

